# Influence of walking speed on locomotor time production

**DOI:** 10.1101/001156

**Authors:** Fabrice Megrot, Carole Megrot

**Affiliations:** Clinical Unit of Gait and Movement Analysis, Physical Medicine and Rehabilitation Center of Bois-Larris, French Red Cross

## Abstract

The aim of the present study was to determine whether or not walking speed affects temporal perception. It was hypothesized that fast walking would reduce the perceived length of time while slow walking increase production estimates. 16 healthy subjects were included. After a first « calibration » phase allowing the determination of different walking speeds, the subjects were instructed to demonstrate periods of time or « target times » of 3s and 7s, by a walking movement. Then, subjects were asked to simulate walking by raising one foot after the other without advancing. Finally, a third condition, Motionless, involved producing the target times while standing without movement. The results of this study suggest that movement does influence the perception of time, causing an overestimation of time. In agreement with the results of Denner et al. (1963) the subjects produced times which were longer than the target times.

---

Temporal information is necessary for interaction with the environment. Humans are capable of perceiving time without the use of instruments. Wearden (2004) showed that subjects were able to correctly compare intervals of time with a standard déviation of 4s. In a review of many experiments, Eisler and al. (1992) found a perfect correlation between target times and times produced by subjects for a range of target times between half a second and a few minutes.

The mechanisms which enable us to evaluate the duration of events within the environment as those of our own movements remain, however, mainly unknown. Although human beings have many receptors, for example the retinas make it possible to perceive light and the organs of Corti make it possible to transform the vibrations of air into sound, we do not have « temporal receptors » (Brown, 1995).

Treisman (1963) proposed that time is perceived by an internal clock mechanism. He proposed that networks of neurons emit stable and regular pulsations which, once stored, allow time to be evaluated. However, other authors (Michon, 1984, Rammsayer & Lima, 1991) have suggested that there are two mechanisms of time perception. The first is of a « highly perceptive » nature (Michon, 1985), similar to the internal clock mechanism described by Treisman (1963) and is used only for the perception of durations of about 500 ms. The second mechanism is more cognitive and acts for durations greater than 500 ms.

Perception of time could thus depend on the perception of events in the environment (Fraisse, 1974, Block & Zackay, 2000) as well as events related to the body, in particular the perception of muscle contractions. Thus, in 1965, Gooddy proposed that the rhythmic movements of our limbs such as occur during walking could be used for the perception of time. Using a « finger-tapping » task in which subjects were asked to tap at different speeds, each for a duration of 70s, Denner et al. (1963) showed that the speed with which the task was carried out influenced the perception of time. When the tapping was slow, the subjects estimated longer time periods than when they tapped quickly. The authors thus made the assumption that the change of movement speed altered the duration perceived. However, the subjects tended to over-estimate the 70 seconds duration, independently of the tapping speed. During the experiments carried out by Newman (1972-1976), the subjects in agreement with Vireordt’s law (Vireordt, 1868) underestimated the 40 seconds duration. Newman (1972) found contrasting results with those of Denner et al. (1963). Newman’s task consisted of producing gait on a treadmill for a duration of 40 seconds. Walking speed was modified by accelerating or slowing down the treadmill. No significant effect of walking speed on the production of the target time of 40s was found, except for a tendency at slow speeds. In further experiments to evaluate the effect of slow walking speeds, Newman (1976) found a significant influence on time perception. The subjects produced longer durations for slower walking speeds. Newman (1976) thus proposed that time perception is affected by slow movement while errors of time perception are compensated for during fast movements. However, during these two experiments, the subjects walked at the speed imposed by the treadmill. They did not actively choose their walking speed. It is thus also possible that time perception is only perturbed when the speed of the movements is actively chosen by the subject. Moreover in another experiment where the subjects were fully actors of their movements, Vercruyssen et al. (1989) demonstrated a significant decrease in the mean value of judged durations during cycling on an ergometer compared to a relaxed phase. The authors suggested that the change in body temperature caused by the cycling effort influenced several timing mechanisms in the central nervous system (Hancock, 1983). But, they did not exclude the possibility that the results may be due to the selected cycling speed of the lower limbs.

The aim of the present study was to determine whether or not walking speed affects temporal perception. It was hypothesized that fast walking would reduce the perceived length of time while slow walking increase production estimates.

If this hypothesis is true then active, unassisted gait would be a factor which influences time perception. The fast or slow execution of movement would generate a poor estimation of time remaining to answer or to perform of an activity.

## Method

### Sample

16 healthy subjects were included, 7 females and 9 males, aged between 21 and 28 years (mean = 23.8, SD = 1.6). All the subjects were volunteers and gave written consent for their participation. Subjects were naive regarding the purpose of the experiment.

None had known perceptual or motor disabilities or took regular drug therapy. Stimulants, depressants, and hallucinogens are known to affect time perception (Meck, 1996).

### Apparatus

An ecological setting was used for the walking conditions. Subjects thus had all their senses such as vision or hearing available.

A VICON MX motion capture system was used to record gait in three dimensions. Subjects wore a black suit with 37 reflective markers positioned on various parts of their body (head, left and right shoulder, sternum, left and right arms, left and right hands, back, left and right hips, left and right thighs, left and right knees, left and right tibias, left and right ankles, left and right toes, and left and right heels). Twelve infrared VICON MX13 cameras were placed along a 25 meter runway creating a 14 meter gait capture zone.

Gait was recorded at a sampling rate of 120 Hz. Temporal data such as stride time and gait velocity were accurate to 0.01 second and spatial data such as stride lengths and the distance covered were accurate to 1 millimeter. The recordings also allowed computation of the global number of steps for each trial.

### Procedure

The entire procedure for this experiment took approximately one hour per subject. Precautions were taken so that the environmental stimuli were the same for all subjects. Also, the lighting and temperature in the testing area were held constant during the experiment because these factors can create variations of time perception (Wearden & Penton-Voak, 1995; Aschoff & Daan, 1997). The experimenter was the only person in the testing area. He maintained the same position throughout the experiment and always read the same specific instructions to the subjects.

After donning the black suit with the reflective markers, the subjects were asked to walk on the runway in the most spontaneous and comfortable manner. The adopted walking speed was then accepted as that subject’s Normal walking speed. The subjects were also requested to walk faster and then slower their Normal walking speed. These spontaneously chosen speeds by the subject were respectively named Fast walking speed and Slow walking speed.

After this first « calibration » phase allowing the determination of the different speeds, the subjects were positioned at the beginning of the runway and asked to carry out a locomotor time production task. The subjects were instructed to demonstrate periods of time or « target times » of 3s and 7s, by a walking movement. The instruction was : « *When you feel ready to begin, walk straight ahead; then when you estimate having walked for the target time, stop precisely at this moment, by joining your two feet* ».

The subjects then carried out this task at their three walking speeds and for both target times; 3s and 7s. The duration of the locomotor time productions was measured between the initiation of the first step and the end of the last step, at the moment when the two feet were joined.

The second condition was On the Spot walking. Subjects were asked to simulate walking by raising one foot after the other without advancing. The third condition, Motionless, involved producing the target times while standing without movement. Subjects raised their right hands to begin the time productions and again to finish it.

Subjects had no feedback on their performance. 5 trials were carried out for each experimental condition, presented in a random order and counterbalanced across subjects.

No limitations or restrictions were placed on the subjects concerning their methods of delimiting the time periods. However, after the experiment, subjects were interviewed regarding their perception of their performance.

### Data analysis

The mean and the standard deviation of produced times per target time, and the mean and standard deviation of the errors of produced time were calculated from the 5 trials carried out by the 16 subjects for each experimental condition (Motionless, On the Spot, Slow, Normal and Fast) and for each target time (3s and 7s).

Errors of produced time were calculated as the time produced by a subject minus the target time. Inferential statistics were used to analyse these different measures; the 2 targets times were evaluated using a 2×5 repeated measures analysis of variance (ANOVA). All post-hoc analyses were carried out using Scheffe’s test, chosen because it is the most conservative of post hoc tests (Winer et al., 1971).

Several gait parameters were also analysed: the mean and standard deviation of the walking rhythm, walking speed and the stride length.

p≤0.05 was considered significant in all cases.

## Results

### Time production

The results obtained for the produced time as a function of the 5 different experimental conditions (Motionless, On the Spot, Slow, Normal and Fast) are presented in Table 1.

**Table 1.**
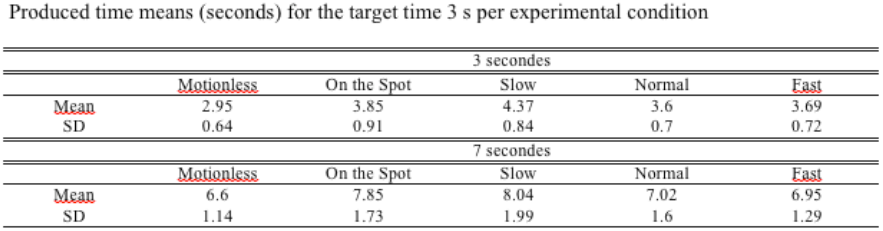
Mean time produced based during the 5 trials for each experimental condition by the 16 subjects; SD, average standard deviations of the 5 time productions carried out by each subject for each experimental condition.

The average times produced for the two target times for the Motionless and On the Spot conditions were higher than those for the other conditions, indicating that subjects tended to overestimate time for these two conditions.

Subjects were more precise for the target time of 7 seconds and errors were greater for the target time of 3 seconds.

The analysis of variance (ANOVA), on the errors produced (F_(4, 250)_ = 15.148, p < 0.001), revealed that the On the Spot (mean = 0.85; SD = 0.97) and slow experimental conditions (mean = 0.39; SD = 1.02), were significantly different from the motionless condition (mean = - 0.23; SD = 0.74). However, there was no difference between the motionless condition and the other experimental conditions (Figure 1).

**Fig. 1.**
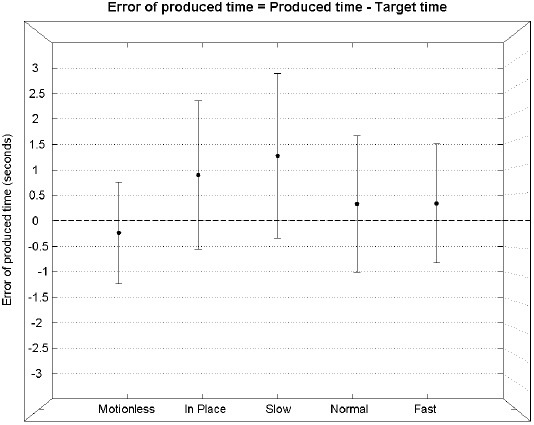
Errors of produced time means as a function of 5 different experimental conditions (Motionless, On the Spot, Slow, Normal and Fast). Error bars represent the maximum and the minimum of each standard deviation.

### Walking analysis

Analysis of the number of steps revealed that the number of walking cycles correlated very strongly with the produced times, r^2^ = 0.69 (p < .001) with a slope of 1.01 (Figure 2).

**Fig. 2.**
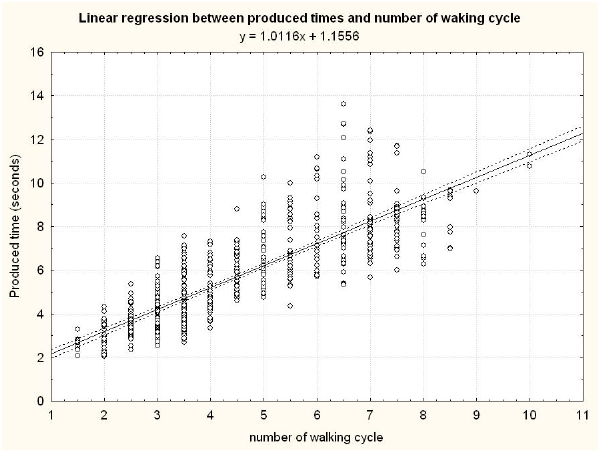
Linear regression between produced times and number of walking cycles. The produced times for the Motionless condition were not taken into account in this linear regression. Dashed lines represent the 95% confidence interval.

Analysis of walking speed (Table 2) showed that the subjects adopted different speeds spontaneously, with an average of 1,66 m/s for fast walking and 0,6 m/s for slow walking, for all the experimental conditions.

**Table 2.** Mean cadence, speed and stride length based on the trials carried out for each experimental condition by the 16 subjects; SD, average standard deviations of the trials carried out by each subject for each experimental condition.

| Cadence |  |  |  |  |  |
| --- | --- | --- | --- | --- | --- |
|  | Motionless | On the Spot | Slow | Normal | Fast |
| Mean | 0.00 | 1.40 | 1.29 | 1.67 | 2.03 |
| SD | 0.00 | 0.27 | 0.16 | 0.14 | 0.19 |
| Walking speed |  |  |  |  |  |
|  | Motionless | On the Spot | Slow | Normal | Fast |
| Mean | 0.00 | 0.00 | 0.62 | 1.00 | 1.44 |
| SD | 0.00 | 0.00 | 0.26 | 0.28 | 0.26 |
| Stride length |  |  |  |  |  |
|  | Motionless | On the Spot | Slow | Normal | Fast |
| Mean | 0.00 | 0.00 | 0.48 | 0.60 | 0.71 |
| SD | 0.00 | 0.00 | 0.17 | 0.13 | 0.10 |

Analysis of the data regarding cadence and stride length (Table 2) showed that the subjects adopted a strategy of reduced cadence and stride length in order to decrease gait velocity and they increased cadence and stride length to increase gait velocity. This conforms with the results of Inman et al. (1981). The subjects thus adopted different cadences for each experimental condition with the fastest rhythm for the Fast experimental condition and similar walking rhythms for the Slow and On the Spot conditions

## Discussion

The results of this study suggest that movement does influence the perception of time, causing an overestimation of time. In agreement with the results of Denner et al. (1963) the subjects produced times which were longer than the target times. However, it is difficult to explain this first result. According to current models of time perception (Zakay and Block, 2004), a dual task (here perception of time and walking), should result in an underestimation of time.

The ANOVA on the errors of produced time also showed that the subjects tended to significantly overproduce time for the On the Spot and Slow experimental conditions. These results are in agreement with those of Newman (1976), suggesting that slow walking has a more significant impact on the perception of time than fast walking. However, the present experiment makes it possible to show that walking speed is not a determining factor of time perception, because subjects also overproduced time in the walking On the Spot condition and there was no significant difference between the Normal and Fast experimental conditions. However, it is necessary to note that the walking speeds of the subjects were relatively slow, 1,66 m/s on average for the Fast condition. Brehmer (1969) showed that for a displacement at 50 Km/h (13,9 m/s), speed had an influence on time perception. In fact, the analysis of the walking rhythms shows that they are slower for the On the Spot and Slow experimental conditions. This may suggest that the slow walking rhythm causes an overproduction of time. The question could thus be raised why a fast cadence does not influence time perception. According to Newman (1976), it is easier to compensate for errors of time perception in the case of overcompensation, i.e. to add time. However, research on the energy cost of walking showed that at slow speeds, oxygen uptake is greater (Inman et al. 1981). This additional effort could thus have a more significant effect on the cognitive processes of the perception of time, causing the overproduction of time in the Motionless and Slow experimental conditions. However, the very strong correlation between the number of gait cycles and times produced show how strongly walking and time perception are interdependent. It is thus probable that the freely selected walking rhythm is a factor influencing the perception of time. Within the framework of the theory developed by Treisman (1963), the overproduction of time for the On the Spot and Slow experimental conditions could be related to a deceleration of the frequency of oscillation of “the internal clock” caused by a slow walking rhythm.

